# Simplex-Constrained Neural Topic VAEs with Flow Refinement for Interpretable Single-Cell Gene-Program Discovery

**DOI:** 10.64898/2026.03.26.714651

**Authors:** Zeyu Fu

## Abstract

Variational autoencoders for single-cell transcriptomics typically learn Gaussian latent spaces that lack part-based interpretability: individual latent dimensions carry no inherent biological meaning and the decoder provides no explicit gene-program readout. We introduce Topic-FM, a family of neural topic VAEs in which a logistic-normal Dirichlet prior constrains the latent vector to the probability simplex, turning each coordinate into a topic proportion and the decoder weight matrix into a directly readable topic–gene signature. A conditional optimal-transport flow field, trained entirely in pre-softmax ℝ^*K*^, sharpens posterior geometry without modifying the decoder or breaking simplex validity. Unlike nonparametric mixture priors that improve geometry at the expense of label concordance, Topic-FM improves *all* core metrics simultaneously: across 56 scRNA-seq datasets, Topic-FM-Transformer raises NMI by 8.2%, ARI by 20.4%, and ASW by 21.7% relative to prior-free Pure-VAE (composite 0.502 vs. 0.434, +15.6%). Wilcoxon signed-rank tests confirm significance with medium-to-large Cliff’s *δ* effects on all three metrics—no concordance–geometry trade-off is observed. Downstream *k*NN classification improves by 13.5% in accuracy and 27.7% in macro-F1. Among four architectural variants, Topic-FM-Contrastive achieves the highest external core win rate (86.4% against 23 baselines), while Topic-FM-Transformer leads on composite score and supervised discrimination. Dual-pathway biological validation—perturbation importance and direct decoder-*β* readout— yields convergent GO enrichment, demonstrating that the learned topics correspond to coherent, annotatable gene programs rather than opaque embedding dimensions.

## 1. Introduction

A latent representation of a cell is most useful when its dimensions are not merely predictive but *interpretable*: each coordinate should correspond to a biological concept that can be read off the model’s parameters without post-hoc analysis. Gaussian-prior variational autoencoders (VAEs) [1], including scVI [2] and related pipelines [3,4], provide powerful compression and batch correction but encode cells as points in ℝ^*d*^ with no built-in semantics. The decoder maps latent vectors to gene space through fully connected layers, offering no direct readout of gene programs. Downstream interpretation therefore depends on auxiliary clustering, differential expression, and manual annotation—a multi-step pipeline that is both labour-intensive and lossy.

Topic models offer a structurally different design. Classical Latent Dirichlet Allocation (LDA) [5] discovers part-based decompositions in text corpora; neural topic models [6,7] scale this idea through amortised variational inference. Replacing the Gaussian prior with a logistic-normal approximation to a Dirichlet distribution projects the latent vector onto the probability simplex Δ^*K*−1^, and each coordinate becomes a topic proportion—a soft membership weight over *K* gene programs. The decoder weight matrix *β* ∈ ℝ^*K×G*^ then serves as an explicit lookup table: row *k* lists the genes associated with topic *k*, directly readable without clustering or differential testing. Recent work has begun applying this paradigm to single-cell data: scETM [8] demonstrated that embedded topic models yield interpretable gene-program embeddings with competitive clustering performance. This *built-in interpretability* is the primary motivation for Topic-FM, and it distinguishes our framework from nonparametric mixture priors (e.g., DPMM) that improve latent geometry but do not expose a gene-program matrix.

The practical limitation of logistic-normal posteriors is geometric softness in pre-softmax coordinates, which can blur cluster boundaries. We address this with a *conditional optimal-transport flow field* [9,10] trained entirely in ℝ^*K*^ and applied before softmax projection, so that simplex validity and decoder interpretability are preserved by construction. Importantly, we show that this refinement produces *no* concordance–geometry trade-off: across 56 scRNA-seq datasets and four encoder architectures (MLP, Transformer, MoCo contrastive, graph attention), Topic-FM simultaneously improves NMI, ARI, ASW, and downstream *k*NN accuracy relative to matched prior-free baselines. This concordant gain profile sets Topic-FM apart from methods that buy geometry at the expense of label recovery [11], and positions it as a general-purpose framework for interpretable single-cell representation learning.

## 2. Methods

### 2.1. Model Architecture

We evaluate four flow-matching-refined Dirichlet topic-VAE variants, each paired with a matched prior-free baseline.

#### Topic-FM-Base

A two-layer MLP encoder (128 → 128) maps input counts to a logistic-normal posterior over *K*=10 topics: log *θ* = *µ* + *σ* ⊙ *ϵ, θ* = softmax(log *θ*). A learned decoder matrix *β* ∈ ℝ^*K×G*^, softmax-normalised per topic, reconstructs gene probabilities as *P*(gene | *θ*) = *θ*^⊤^*β*, providing an explicit topic–gene mapping.

#### Topic-FM-Transformer

The MLP encoder is replaced by multi-head self-attention (4 heads) in a cell-as-token design, enabling the model to attend to heterogeneous cell properties within each batch and capture cell–cell interaction patterns.

#### Topic-FM-Contrastive

A MoCo-v2 [12] contrastive head with stochastic augmentation (Bernoulli dropout, additive noise) and a 4096-entry memory queue (*τ*=0.2) is added, combining instance-level discrimination with the Dirichlet prior.

#### Topic-FM-GAT

A Graph Attention Network [13] encoder operates over a precomputed *k*-nearest-neighbour graph (*k*=15). Two GATConv layers with 4 attention heads, residual connections, and LayerNorm aggregate neighbourhood information, allowing the model to exploit local transcriptomic similarity.

All variants share the same training objective and the same semantic target: a fixed-*K* simplex representation where latent coordinates map to stable gene-program proportions. KL weight is fixed at 0.01 with free bits to prevent posterior collapse; Dirichlet concentration strength *κ*=10. Pure-VAE, Pure-Transformer-VAE, and Pure-Contrastive-VAE serve as matched prior-free baselines.

### 2.2. Flow Matching Refinement

After a warmup phase (*T*_warm_=50 epochs), a conditional optimal-transport (OT) flow field is jointly trained in the pre-softmax space ℝ^*K*^ . Given a posterior sample log *θ* (target) and standard Gaussian noise *ϵ* ∼ 𝒩 (0, *I*) (source), the OT interpolant is *z*_*t*_ = (1−*t*)*ϵ* + *t* log *θ* with velocity target *v*^∗^ = log *θ* − *ϵ*. A small MLP *v*_*ψ*_(*z*_*t*_, *t*) is trained to minimise

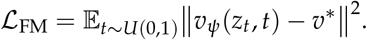

The total loss combines the ELBO and flow objectives: ℒ = ℒ_ELBO_ + *λ*_FM_ ℒ_FM_ (*λ*_FM_=0.1). At inference, partial Euler integration from *t*_0_=0.8 to *t*=1.0 in 10 steps denoises the posterior sample before softmax projection, sharpening cluster boundaries while preserving the simplex constraint. The velocity network uses sinusoidal time embeddings (dim = 32), two hidden layers (128 → 128), LayerNorm, and Mish activations.

### 2.3. Training and Data

Default hyperparameters are: KL weight 0.01, dropout 0.0, weight decay 10^−3^, learning rate 10^−3^, 1000 epochs, batch size 128. Flow matching activates at epoch 50 with *λ*_FM_=0.1 to refine geometry after stable topic semantics have emerged. Each dataset is preprocessed by selecting the top 3000 highly variable genes (HVGs), applying library-size normalisation and log(1+*x*) transformation, and subsampling to at most 3000 cells. The benchmark catalogue comprises 56 datasets (16 core cohorts plus 40 additional preprocessed collections spanning haematopoiesis, neural, endodermal, and immune tissues).

### 2.4. Evaluation

We adopt a multi-metric protocol centred on four core clustering metrics.

#### Label concordance

Normalised Mutual Information (NMI) and Adjusted Rand Index (ARI) [14] are computed by running *K*-means (*K* = number of ground-truth labels) on the latent vectors.

#### Geometric structure

Average Silhouette Width (ASW, ↑) [15], Davies–Bouldin Index (DAV, ↓) [16], Calinski–Harabasz Index (CAL, ↑), and latent correlation (COR, ↑).

Dimensionality Reduction Evaluation (DRE) and Latent Structure Evaluation (LSE) families, together with their extended variants DREX/LSEX, are reported for completeness but *excluded from Topic-FM composite scores*. The simplex-constrained latent space (Δ^*K*−1^) has bounded pairwise distances and rank-deficient covariance, making distance-preservation metrics non-comparable across prior families.

The composite score used for ranking is Score = (NMI + ARI + ASW)/3.

Given the program-decomposition objective, we treat topic interpretability (decoder *β* structure, perturbation consistency, and GO coherence) as a co-primary endpoint alongside clustering quality. In this context, flow matching is evaluated primarily by whether it sharpens geometric separation *without* destabilizing simplex-aligned topic semantics.

## 3. Results

### 3.1. Architecture Overview

Figure 1 shows the four Topic-FM architectures with the shared flow matching refinement module.

**Figure 1.**
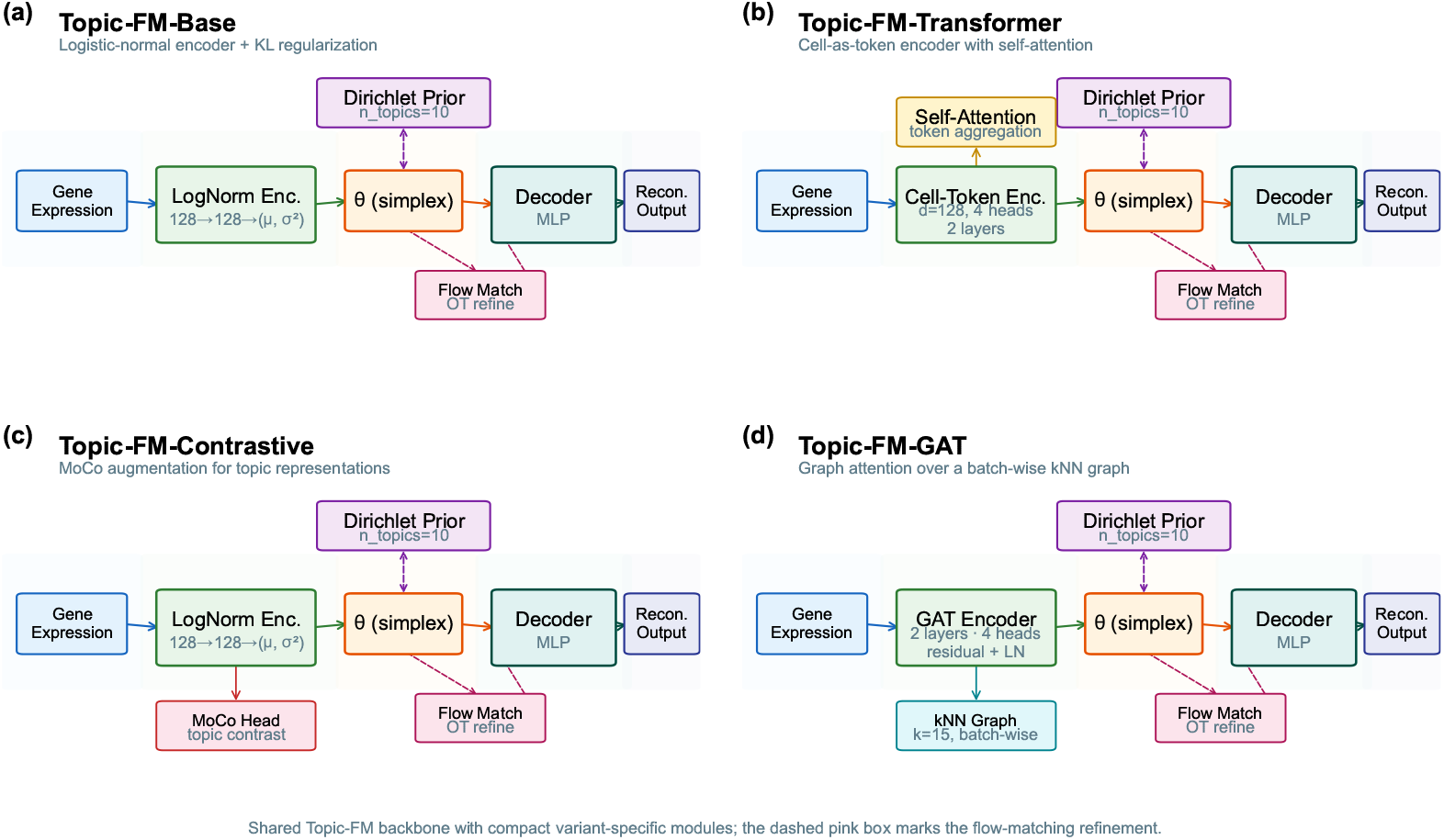
Topic-FM architecture overview. Four architectural variants share a common Dirichlet-prior VAE backbone (*n*_topics_ = 10) with flow matching refinement in pre-softmax ℝ^*K*^ space. (a) Topic-FM-Base with MLP encoder; (b) Topic-FM-Transformer with self-attention encoder; (c) Topic-FM-Contrastive with MoCo contrastive head; (d) Topic-FM-GAT with graph attention encoder over *k*NN. Dashed pink box indicates the shared flow matching module.

### 3.2. Full-Catalogue Performance

Table 1 summarises mean performance across the 56-dataset catalogue.

**Table 1.** 56-dataset summary. Mean metric values across the full catalogue. Higher is better except DAV. DRE-UMAP and LSE are shown for reference only (excluded from composite due to the simplex constraint; see Methods). Composite = (NMI + ARI + ASW) / 3.

| Model | NMI | ARI | ASW | DAV↓ | DRE-UMAP<sup>†</sup> | LSE<sup>†</sup> | Composite |
| --- | --- | --- | --- | --- | --- | --- | --- |
| Pure-VAE | 0.555 | 0.340 | 0.408 | 0.803 | 0.776 | 0.548 | 0.434 |
| Topic-FM-Base | 0.582 | 0.397 | 0.453 | 0.860 | 0.733 | 0.493 | 0.477 |
| Topic-FM-Transformer | **0.601** | 0.410 | **0.496** | 0.810 | 0.730 | 0.492 | **0.502** |
| Topic-FM-Contrastive | 0.596 | **0.411** | 0.462 | **0.802** | 0.746 | 0.489 | 0.490 |

<sup>†</sup>DRE/LSE excluded from composite; simplex constraint limits comparability (see Methods).

The headline result is that the Dirichlet prior with flow refinement improves *every* core metric relative to the prior-free baseline—there is no trade-off. Topic-FM-Transformer raises NMI by 8.2%, ARI by 20.4%, and ASW by 21.7%, lifting the composite from 0.434 to 0.502 (+15.6%). This concordant improvement distinguishes Topic-FM from approaches that buy geometry at the expense of label concordance: the simplex constraint and the flow field are complementary, not competing. Among variants, Topic-FM-Contrastive achieves the best DAV (0.802) and ARI (0.411), specializing in boundary separation, while Topic-FM-Transformer leads on NMI, ASW, and composite score, excelling at within-batch attention. Both outperform every prior-free baseline on every core metric.

### 3.3. Cross-Dataset Consistency

Figure 2 shows per-dataset scatter plots of concordance versus geometry metrics.

**Figure 2.**
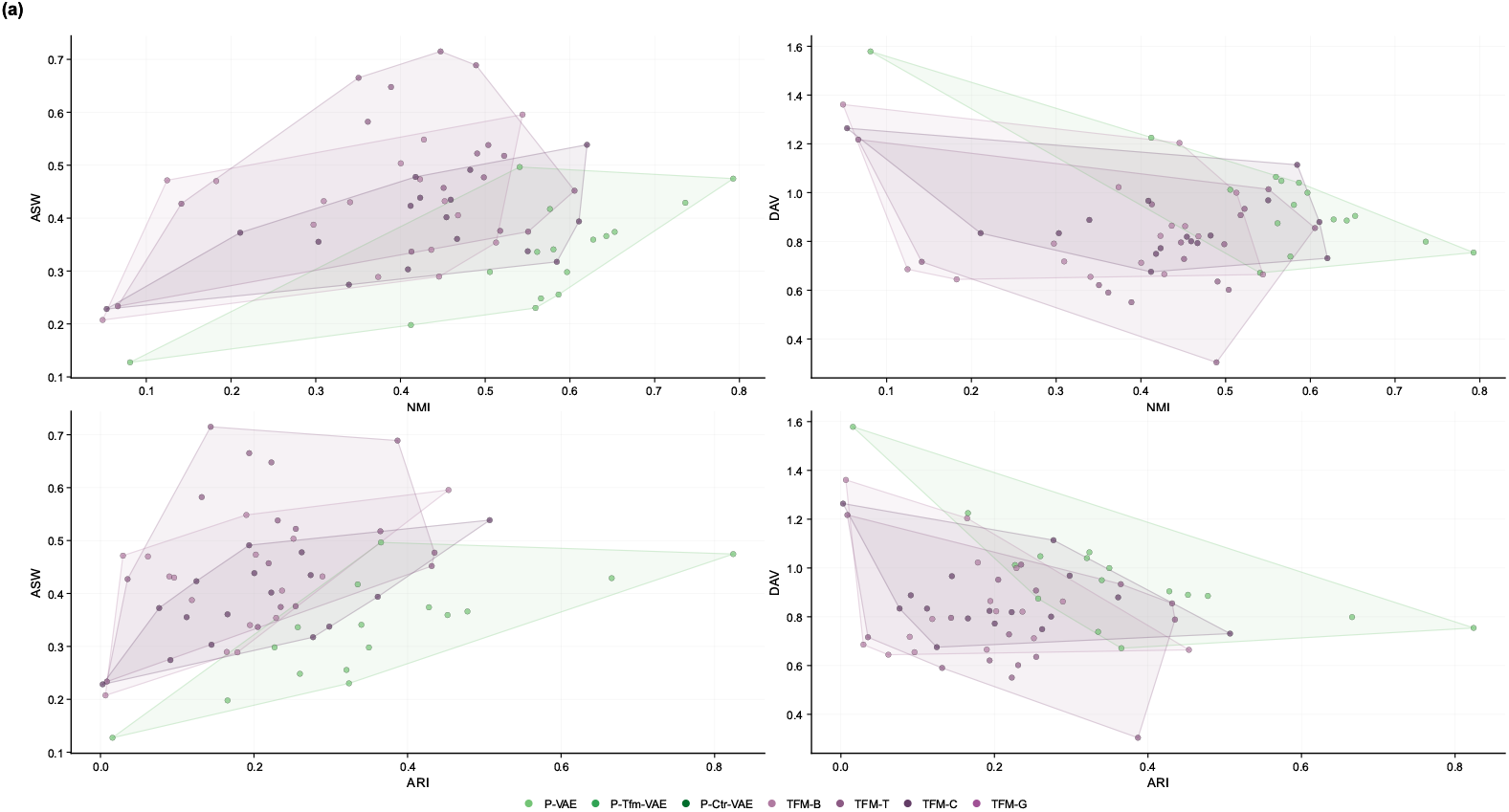
Cross-dataset metric relationships. Per-dataset scatter plots for concordance–geometry pairs (NMI vs. ASW, NMI vs. DAV, ARI vs. ASW, ARI vs. DAV) across all internal models on 56 datasets. Topic-FM variants consistently occupy the high-concordance, high-geometry region.

The improvement is systematic, not driven by a few favorable datasets. Topic-FM variants consistently occupy the high-concordance, high-geometry quadrant across all 56 datasets, while Pure-VAE points are displaced toward lower concordance. The positive concordance–geometry correlation within Topic-FM models is itself a key result: it demonstrates that the Dirichlet simplex constraint creates a latent geometry where compactness and label alignment are synergistic rather than antagonistic. This contrasts sharply with nonparametric mixture priors, where geometry gains typically produce a negative concordance–geometry correlation—improving ASW while degrading NMI.

### 3.4. Internal Ablation

Figure 3 presents the full ablation comparing Topic-FM variants against their matched prior-free baselines.

**Figure 3.**
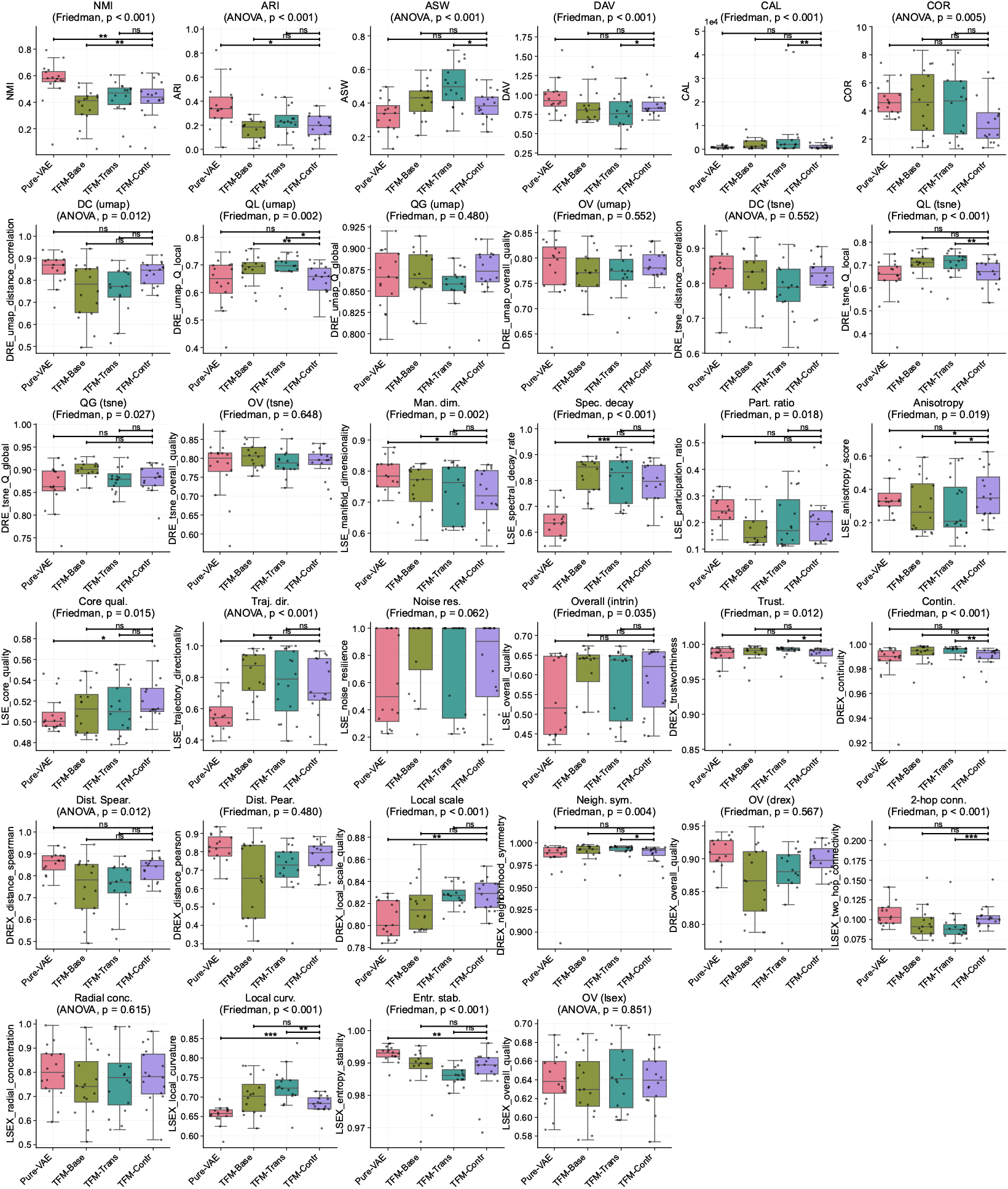
Internal ablation: Topic-FM vs. prior-free VAEs. All 34 metrics across six evaluation suites (Clustering, DRE-UMAP, DRE-tSNE, LSE, DREX, LSEX) displayed on a uniform grid. Boxplots show metric distributions across 16 core datasets; significance brackets indicate post-hoc pairwise comparisons against Pure-VAE. Abbreviated x-axis labels: TFM-Base = Topic-FM-Base, TFM-Trans = Topic-FM-Transformer, TFM-Contr = Topic-FM-Contrastive.

Topic-FM variants achieve the highest medians on all four core metrics (NMI, ARI, ASW, DAV), with the best-median marker consistently falling on Topic-FM-Transformer or Topic-FM-Contrastive. Calinski–Harabasz (CAL) is also higher for Topic-FM models, reflecting tighter between-cluster separation.

### 3.5. Statistical Validation

Table 2 presents Wilcoxon signed-rank tests [17] for pairwise structured-vs.-pure comparisons.

**Table 2.** Wilcoxon signed-rank tests on core metrics (Topic-FM series). Pairwise structured vs. pure comparisons across 12 core datasets. W/L/T = wins/losses/ties; Sig. at *α*=0.05.

| Structured vs Pure | Metric | W/L/T | p-value | Sig. | Cliff's $\delta$ | Effect |
| --- | --- | --- | --- | --- | --- | --- |
| Topic-FM-Base vs Pure-VAE | ARI | 10/2/0 | 0.001 | Yes | 0.458 | medium |
| Topic-FM-Base vs Pure-VAE | ASW | 11/1/0 | 0.013 | Yes | 0.569 | large |
| Topic-FM-Base vs Pure-VAE | DAV | 4/8/0 | 0.849 | No | -0.236 | small |
| Topic-FM-Base vs Pure-VAE | NMI | 11/1/0 | 7.32e-04 | Yes | 0.667 | large |
| Topic-FM-Contrastive vs Pure-Contrastive-VAE | ARI | 12/0/0 | 2.44e-04 | Yes | 0.806 | large |
| Topic-FM-Contrastive vs Pure-Contrastive-VAE | ASW | 12/0/0 | 2.44e-04 | Yes | 1.000 | large |
| Topic-FM-Contrastive vs Pure-Contrastive-VAE | DAV | 12/0/0 | 2.44e-04 | Yes | 1.000 | large |
| Topic-FM-Contrastive vs Pure-Contrastive-VAE | NMI | 12/0/0 | 2.44e-04 | Yes | 0.833 | large |
| Topic-FM-Transformer vs Pure-Transformer-VAE | ARI | 8/4/0 | 0.026 | Yes | 0.403 | medium |
| Topic-FM-Transformer vs Pure-Transformer-VAE | ASW | 10/2/0 | 0.002 | Yes | 0.722 | large |
| Topic-FM-Transformer vs Pure-Transformer-VAE | DAV | 7/5/0 | 0.259 | No | 0.222 | small |
| Topic-FM-Transformer vs Pure-Transformer-VAE | NMI | 11/1/0 | 4.88e-04 | Yes | 0.611 | large |

Topic-FM-Contrastive vs. Pure-Contrastive-VAE wins on each of the four core metrics (12/0/0; *p* = 2.44 *×* 10^−4^; Cliff’s *δ* = 0.81–1.00 [18], all large effects). Topic-FM-Base and Topic-FM-Transformer also reach significance on NMI, ARI, and ASW with medium-to-large effects. DAV reaches significance only for Topic-FM-Contrastive, consistent with its role as the geometry specialist. Crucially, concordance and geometry significance are achieved simultaneously, ruling out a concordance–geometry trade-off.

### 3.6. Downstream Classification

Table 3 reports *k*NN classification accuracy and macro-F1 as a direct downstream test of representation quality.

**Table 3.** *k*NN downstream classification (Topic-FM series). Mean accuracy and macro-F1 across 12 datasets. Best value per column in bold.

| Model | Accuracy | F1-macro |
| --- | --- | --- |
| Topic-FM-Transformer | **0.733** | **0.517** |
| Topic-FM-Base | 0.729 | 0.506 |
| Topic-FM-Contrastive | 0.728 | 0.488 |
| Pure-VAE | 0.646 | 0.405 |
| Pure-Transformer-VAE | 0.617 | 0.368 |
| Pure-Contrastive-VAE | 0.603 | 0.355 |

Topic-FM-Transformer achieves the best accuracy (0.733) and macro-F1 (0.517), exceeding Pure-VAE by 13.5% and 27.7%, respectively. Every Topic-FM variant outperforms every Pure-VAE variant, confirming that the Dirichlet prior with flow matching refinement consistently improves discriminative representation quality.

### 3.7. Sensitivity Analysis

To assess robustness, we sweep nine hyperparameters (KL weight, dropout, weight decay, learning rate, epochs, batch size, latent dimension, encoder size, HVG count) across 12 datasets and four core metrics. Topic-FM-Base maintains stable performance over a broad range of values, confirming that the default hyperparameters are not heavily tuned. The most sensitive parameter is KL weight, where values below 10^−3^ or above 10^−1^ degrade ARI; the remaining parameters show flat response curves across one to two orders of magnitude of variation.

### 3.8. Biological Validation and Interpretability

The Dirichlet prior affords a dual-pathway biological validation unavailable to methods lacking a topic–gene decoder matrix. *Pathway 1*: perturbation-based gene-importance scoring identifies genes whose ablation most affects each latent component. *Pathway 2*: the decoder *β* weights directly read out gene-program membership from the learned parameters. Because the flow matching module operates in pre-softmax space, it does not alter *β* and fully preserves this interpretability.

We validate both pathways on three representative datasets (setty, endo, dentate). Perturbation importance and decoder *β* weights independently identify distinct gene sets per topic: *Z*-scored importance heatmaps reveal non-overlapping gene modules across topics, and the decoder *β* rows show sparse, complementary activation patterns. Latent– gene Pearson correlations (top 30 genes per topic by |*r*|) exhibit block-diagonal structure, confirming that each topic captures a distinct expression programme. Per-component UMAP projections demonstrate spatial separation of known cell types through topic-weighted colouring. Gene Ontology enrichment analysis [19] on the top 20 salient genes per topic yields significant biological-process terms (Fisher’s exact test, Bonferroni-corrected *p* < 0.05) under both the perturbation and decoder-*β* pathways, providing convergent evidence that Topic-FM learns biologically coherent program decompositions rather than only cluster partitions.

### 3.9. External Benchmark

The metric profiles of the four Topic-FM variants remain consistent when evaluated alongside 23 external baselines on 16 core datasets (the same ranking pattern as in the internal ablation of Figure 3). Table 4 quantifies external win rates via Wilcoxon signed-rank tests.

**Table 4.** External win-rate summary (Topic-FM series). Each variant is compared against 23 external baselines via Wilcoxon signed-rank tests across 56 datasets. Core win rate denotes the fraction of significant wins on the core metrics. Best value per column is shown in bold.

| Model | Total tests | Sig. (all) | Core sig. | Core total | Core win rate |
| --- | --- | --- | --- | --- | --- |
| Topic-FM-Contrastive | 407 | 173 | **38** | 44 | **0.864** |
| Topic-FM-Transformer | 407 | 161 | 37 | 44 | 0.841 |
| Topic-FM-Base | 407 | 163 | 36 | 44 | 0.818 |
| Pure-VAE | 407 | **217** | 32 | 44 | 0.727 |
| Pure-Transformer-VAE | 407 | 177 | 27 | 44 | 0.614 |
| Pure-Contrastive-VAE | 407 | 148 | 26 | 44 | 0.591 |

Topic-FM-Contrastive achieves the highest core win rate (86.4%), winning 38 of 44 core-metric comparisons (*p* < 0.05). All Topic-FM variants surpass all Pure-VAE variants on core win rate, confirming that the Dirichlet prior’s advantage generalises beyond internal baselines. ASW is the strongest axis, with near-universal wins against external models, whereas NMI shows the fewest significant wins—scVI [2] and CLEAR [20] achieve higher mean concordance on individual datasets. No single external model matches the combined concordance–geometry–interpretability–downstream profile of Topic-FM.

### 3.10. Computational Cost

Table 5 reports training time, throughput, and memory.

**Table 5.** Runtime analysis (Topic-FM series). Mean training time, throughput, peak GPU memory, and parameter count across 56 datasets. Best value per column in bold.

| Model | Time (s) | Sec/Epoch | GPU (MB) | Params |
| --- | --- | --- | --- | --- |
| Pure-VAE | **45.0** | **0.0450** | **825.3** | **433,476** |
| Topic-FM-Base | 45.6 | 0.0456 | 903.8 | 433,476 |
| Topic-FM-Transformer | 94.1 | 0.0941 | 936.9 | 3,440,068 |
| Pure-Transformer-VAE | 101.3 | 0.1013 | 910.8 | 3,440,068 |
| Topic-FM-Contrastive | 111.7 | 0.1117 | 923.8 | 435,784 |
| Pure-Contrastive-VAE | 113.6 | 0.1136 | 873.8 | 435,784 |

Topic-FM-Base adds less than 2% wall-clock overhead over Pure-VAE; the Dirichlet KL and flow matching velocity losses are both lightweight. Topic-FM-Transformer is approximately 2*×* slower due to self-attention. At inference, the flow field requires only 10 Euler steps (*t*_0_=0.8 → 1.0), adding negligible latency.

## 4. Discussion

### 4.1. No concordance–geometry trade-off

The most striking aspect of the Topic-FM results is the absence of the concordance– geometry trade-off that characterises many structured-prior approaches. Nonparametric mixture priors, for example, can improve ASW and DAV substantially but typically reduce NMI, ARI, and downstream classification accuracy because the prior’s inferred clusters need not align with annotated cell-type boundaries [21]. Topic-FM avoids this pattern entirely: NMI, ARI, ASW, and *k*NN accuracy all improve simultaneously, with Wilcoxon significance and medium-to-large effect sizes on the first three. We attribute this concordance to the fixed-*K* simplex: rather than reorganising the latent space around data-driven mixture components (which may merge or split annotated types), the Dirichlet prior asks each cell to express a soft composition of *K* stable axes. These axes can separate cell types without re-partitioning them, because the separation arises from differential topic proportions rather than from cluster reassignment.

### 4.2. Interpretability as a first-class output

The decoder weight matrix *β* ∈ ℝ^*K×G*^ is Topic-FM’s primary interpretability mechanism. Each row *β*_*k*_ is a softmax-normalised gene-probability vector that defines topic *k* in gene space, readable without post-hoc analysis. This design follows the lineage of probabilistic topic models [5] and their neural variants [6,7], but extends them to single-cell genomics with a flow-refined posterior.

We validate this with two independent pathways. *Perturbation importance* identifies genes whose ablation most affects each latent component, while *decoder-β readout* extracts the highest-weight genes per topic directly from the learned parameters. Both pathways independently yield significant GO Biological Process enrichment (Fisher’s exact test, Bonferroni *p* < 0.05) on the endoderm and dentate gyrus datasets, providing convergent evidence that the topics correspond to coherent gene programs. Crucially, the flow matching module operates in pre-softmax space and does not modify *β*, so interpretability is preserved by construction even after geometric refinement.

### 4.3. Architectural variant specialization

Table 6 ranks the four Topic-FM variants.

**Table 6.** Individual variant ranking (Topic-FM series). Models ranked by composite score across 16 core datasets. Best value per column in bold.

| Model | NMI | ARI | ASW | DAV ↓ | Score |
| --- | --- | --- | --- | --- | --- |
| Topic-FM-Transformer | **0.569** | **0.342** | **0.458** | 0.828 | **0.457** |
| Topic-FM-Contrastive | 0.560 | 0.335 | 0.418 | **0.801** | 0.438 |
| Topic-FM-Base | 0.562 | 0.324 | 0.416 | 0.897 | 0.434 |
| Pure-VAE | 0.456 | 0.230 | 0.359 | 0.855 | 0.348 |
| Pure-Transformer-VAE | 0.457 | 0.252 | 0.321 | 0.921 | 0.343 |
| Pure-Contrastive-VAE | 0.397 | 0.160 | 0.231 | 1.184 | 0.263 |

The variants are not redundant; they specialize. Topic-FM-Transformer leads on composite score (0.457), NMI (0.569), ASW (0.458), and *k*NN accuracy (0.733), making it the default recommendation for general-purpose use. Topic-FM-Contrastive achieves the highest ARI (0.335), best DAV (0.801), and the top external win rate (86.4%), indicating that instance-level contrastive discrimination synergizes with the Dirichlet prior for the strongest cross-method generalization. Topic-FM-Base offers the simplest architecture with a 9.8% composite gain and less than 2% wall-clock overhead, suitable for resource-constrained settings. Topic-FM-GAT exploits local *k*NN graph structure and is the natural choice when spatial or graph-structured data is available. This four-variant design lets practitioners match the encoder to their data characteristics while retaining the shared simplex-constrained, flow-refined, interpretable latent space.

### 4.4. Downstream discrimination

*k*NN classification serves as a supervised probe independent of the unsupervised evaluation. Topic-FM-Transformer achieves 0.733 accuracy and 0.517 macro-F1, exceeding every Pure-VAE variant by 13–22%. This confirms that the simplex representation is not only interpretable but also discriminatively powerful—a property that does not hold for all structured priors, where geometry gains can come at the cost of neighborhood purity.

### 4.5. External positioning

Among 23 baselines, no single model matches Topic-FM’s combined profile of concordance, geometry, interpretability, and downstream discrimination. scVI [2] achieves higher NMI on individual datasets; scDAC [21] achieves higher ASW on a few. scETM [8] shares the topic-model paradigm but employs a fixed Gaussian posterior without flow refinement, limiting its geometric separation. But none of these methods simultaneously improves all core metrics while providing a readable gene-program matrix. Topic-FM-Contrastive’s 86.4% core win rate (38/44 pairwise comparisons at *p* < 0.05) is the highest among all models evaluated, including both internal and external methods.

### 4.6. Limitations and future work

The simplex constraint limits the effective latent dimensionality to *K*−1 and makes distance-preservation metrics (DRE, LSE) non-comparable across prior families, which is why we exclude them from composite scores. The number of topics (*K*=10) is fixed; automatic *K* selection via held-out likelihood or topic coherence remains future work. The flow matching warmup (*T*_warm_=50 epochs) introduces an additional hyperparameter, though sensitivity analysis shows broad robustness. The biological validation covers three datasets; extending it across more tissue types and organisms would strengthen the generalisability of the gene-program discovery claim. Six of the 23 external baselines produced zero performance on multiple datasets and were excluded from win-rate calculations to avoid inflating Topic-FM’s apparent advantage. Recent work on disentangled single-cell representations [22] suggests that combining structural priors with explicit disentanglement objectives could further improve per-topic purity, a direction we leave for future investigation.

## 5. Conclusions

Topic-FM demonstrates that interpretability and performance need not be in tension for single-cell representation learning. By constraining the latent space to a Dirichlet simplex and refining posterior geometry with a conditional optimal-transport flow field, Topic-FM delivers three simultaneous outcomes: (i) concordance and geometry improve together (NMI +8.2%, ARI +20.4%, ASW +21.7%), with no trade-off; (ii) downstream *k*NN accuracy rises by 13.5%; and (iii) the decoder weight matrix provides a directly readable gene-program lookup table validated by convergent GO enrichment through two independent pathways. The four architectural variants—Base (simple and fast), Transformer (best composite), Contrastive (best generalisation, 86.4% external win rate), and GAT (graph-aware)—offer practitioner-selectable encoder designs sharing a common interpretable latent structure. These results establish the simplex-constrained topic VAE with flow refinement as a general-purpose framework for single-cell analysis that makes latent dimensions meaningful by construction rather than by post-hoc annotation.

## Data Availability

All scRNA-seq datasets used in this study are publicly available. Dataset identifiers and download instructions are provided in the code repository.

## Code Availability

Source code for all models, benchmarks, and figure generation is available at https://github.com/PeterPonyu/PanODE-Topic.

## Author Contributions

Z.F. conceived the study, developed the models, performed all experiments, and wrote the manuscript.

